# *Marchantia polymorpha* model reveals conserved infection mechanisms in the vascular wilt fungal pathogen *Fusarium oxysporum*

**DOI:** 10.1101/2021.03.20.436100

**Authors:** Amey Redkar, Selena Gimenez Ibanez, Mugdha Sabale, Bernd Zechmann, Roberto Solano, Antonio Di Pietro

## Abstract

The non-vascular plant *Marchantia polymorpha* has emerged as a valuable model for studying evolutionarily conserved microbial infection strategies and plant immune responses. However, only a handful of fungal pathogens of *Marchantia* have been described so far. Here we establish a new pathosystem using the root-infecting vascular wilt fungus *Fusarium oxysporum*. On angiosperms, this fungus exhibits exquisite adaptation to the plant vascular niche and host-specific pathogenicity, both of which are conferred by lineage-specific effectors secreted during growth in the xylem. We show that *F. oxysporum* isolates with different lifestyles - pathogenic or endophytic - are able to infect this non-vascular liverwort causing tissue maceration and plant cell killing. Similar to bacterial pathogens, *F. oxysporum* induces a PAMP-triggered immune response in *M. polymorpha*. Analysis of isogenic fungal mutants established that infection of *Marchantia* requires conserved fungal pathogenicity mechanisms such as mitogen activated protein kinases, transcriptional regulators and cell wall remodeling enzymes. Remarkably, lineage-specific virulence effectors are dispensable for infection, most likely due to the absence of xylem tissue in this non-vascular plant. The *F. oxysporum* - *M. polymorpha* system provides new insights into the mechanism and evolution of pathogenic and endophytic fungus-plant interactions.

**Significance statement:** Root-infecting vascular fungi cause wilt diseases and provoke devastating losses in hundreds of crops. It is currently unknown how these pathogens evolved and whether they infect non-vascular plants, which diverged from vascular plants over 450 million years ago. Here we show that two strains of the fungus *Fusarium oxysporum* with opposed lifestyles, causing either wilting and death or beneficial protection on tomato, produce similar disease symptoms on the non-vascular plant *Marchantia polymorpha.* We define a set of core fungal pathogenicity factors required on both vascular and non-vascular plants and show that host-specific effectors contributing to disease on tomato are dispensable on *Marchantia*. These findings suggest that systemic wilt disease evolved in fungal pathogens after the emergence of vascular land plants.

## Main Text

### Introduction

How co-evolution has shaped the interaction between plants and their associated microbes remains a central question in organismic interactions (1, 2). Plants have evolved a sophisticated and multi-layered immune system to ward off potential microbial invaders (3–5). Meanwhile, pathogens have developed mechanisms allowing them to enter the living plant, colonize its tissues and overcome its defense responses. Pathogenicity factors can be either broadly conserved or pathogen-specific and include regulators of cell signalling, gene expression or development, as well as secreted effector molecules that modulate the host environment (6–11).

A particularly destructive group of plant root pathogens are those causing vascular wilt diseases, which colonize the highly protected and nutrient poor niche of the xylem (12). The ascomycete fungus *Fusarium oxysporum* (Fo) represents a species complex with worldwide distribution that provokes devastating losses in more than a hundred different crops (13). The fungus is able to locate roots in the soil by sensing secreted plant peroxidases via its sex pheromone receptors and the cell wall integrity mitogen activated protein kinase (MAPK) pathway (14, 15). Inside the root, *F. oxysporum* secretes a small regulatory peptide that mimicks plant <u>R</u>apid <u>AL</u>kalinization <u>F</u>actor (RALF) to induce host alkalization, which in turn activates a conserved MAPK cascade that promotes invasive growth (16). Individual isolates of *F. oxysporum* exhibit host specific pathogenicity, which is determined by lineage specific (LS) genomic regions that encode distinct repertoires of effectors known as Secreted in Xylem (Six) (17, 18). Some Six proteins appear to primarily target plant defense responses, but can also be recognized by specific host receptors (19, 20). Besides the pathogenic forms, the *F. oxysporum* species complex also includes endophytic isolates such as Fo47, which was originally isolated from a natural disease suppressive soil (21). Fo47 colonizes plant roots without causing wilt disease and functions as a biological control agent against pathogenic *F. oxysporum* strains. How vascular fungi such as *F. oxysporum* have evolved and how they switch between endophytic and pathogenic lifestyles remains poorly understood.

The bryophyte *Marchantia polymorpha* belongs to the early-diverged lineage of liverworts and has emerged as the prime non-vascular plant model for addressing the evolution of molecular plant microbe interactions (Evo-MPMI), due to its low genetic redundancy, the simplicity of its gene families and an accessible molecular genetic toolbox (22–26). Importantly, this early plant model possesses receptor-like kinases (RLKs), Nucleotide Binding, Leucine-rich Repeat receptors (NLRs) and salicylic acid (SA) pathway genes which mediate immune signaling in angiosperms (24, 27), allowing the study of plant-microbe interactions across evolutionarily distant land plant lineages, such as liverworts and eudicots, which diverged more than 450 million years ago (28). However, a major shortcoming of *M. polymorpha* is that only few pathogen infection models have been established so far. These include the oomycete *Phytopthora palmivora*, the fungus *Colletotrichum sp1* and the gram-negative bacterium *Pseudomonas syringae* (26, 29–30). A survey of the *M. polymorpha* microbiome identified fungal endophytes that can also act as pathogens (31–32). Whether root infecting vascular fungi can colonize this early-diverging land plant lineage, which lacks both true roots and xylem, is currently unknown.

Here we established a new pathosystem between *F. oxysporum* and *M. polymorpha.* We find that both pathogenic and endophytic *F. oxysporum* isolates can infect, colonize and macerate the thallus of this non-vascular plant. Infection by *F. oxysporum* requires fungal core pathogenicity factors, whereas lineage-specific effectors are dispensable. These results provide new insights into evolutionarily ancient infection mechanisms of vascular wilt pathogens and suggests that the differentiation into endophytic and pathogenic wilting lifestyles evolved after the emergence of vascular land plants.

### Results

#### *F. oxysporum* strains with different lifestyles can infect *M. polymorpha*

To understand the emergence and origin of vascular pathogen infection, we first set out to test whether the non-vascular liverwort *M. polymorpha* can be infected by a root vascular pathogen, which is highly adapted to growth in the xylem of angiosperm plants. Thalli of the accession Tak-1 were inoculated by dipping the abaxial surface in a suspension of microconidia, a common infection protocol for wilt pathogens. *M. polymorpha* thalli inoculated with the tomato pathogenic *F. oxysporum* isolate Fol4287 exhibited disease symptoms including chlorosis and progressive maceration of the thallus tissue, that were absent in the mock treated controls (Fig. 1A and *SI Appendix*, Fig. S1A). The symptoms remained mostly localized to the centre of the mature thallus, while the meristematic apical notches were less affected and often regenerated after 30 to 40 dpi (*SI Appendix*, Fig. S1B). A similar pattern of infection was previously reported for the bacterial pathogen *P. syringae* (26). Symptom severity was dependent on the inoculum concentration (*SI Appendix*, Fig. S1A). The endophytic *F. oxysporum* isolate Fo47 caused similar disease symptoms as Fol4287 (Fig. 1A and *SI Appendix*, Fig. S1B), although tissue maceration progressed slower compared to Fol4287. An alternative drop inoculation method on the surface of the thallus, as previously described for *P. palmivora* (28), resulted in similar symptom development as the dip inoculation protocol, although severe maceration was never observed (*SI Appendix*, Fig. S1C).

**Fig. 1.**
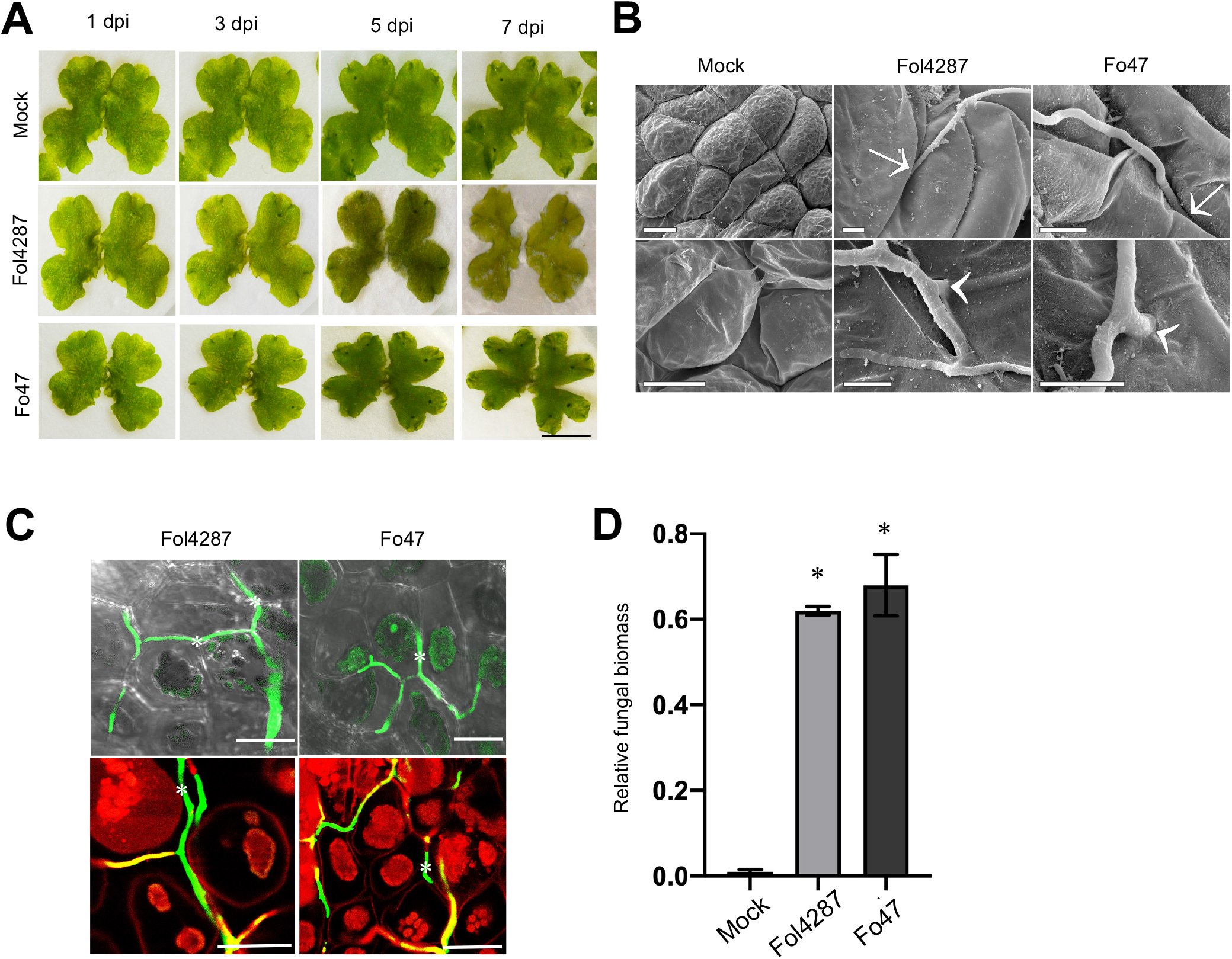
*Fusarium oxysporum* strains with different lifestyles can infect *Marchantia polymorpha*. **(A)** Macroscopic disease symptoms on *M. polymorpha* Tak-1 plants 1, 3, 5 and 7 days after dip inoculation with 10^6^ microconidia ml^−1^ of the *Fusarium oxysporum* (Fo) strains Fol4287 (tomato pathogen) or Fo47 (endophyte), or water (mock). Images are representative of three independent experiments. Scale bar, 1 cm. **(B)** Scanning electron micrographs showing hyphal penetration events on *M. polymorpha* Tak-1 plants 3 days after dip inoculation with the indicated *Fo* strains or water (mock). Arrows = intercellular penetration; arrowheads = intracellular penetration. Scale bar, 20 μm in upper mock image, 5 μm in all others. **(C)** Confocal microscopy demonstrating intercellular hyphal growth of the indicated *Fo* strains expressing clover on TAK1 plants at 3 dpi. Plant cells were stained with propidium iodide (red). Intercellular hyphal growth *in planta* are denoted with an asterisk. Scale bars, 25 μm. **(D)** Quantification of fungal biomass on *M. polymorpha* Tak-1 plants 6 days after dip inoculation with the indicated Fo strains or water (mock). Fungal biomass was measured by real time qPCR using specific primers for the *six1* (Fol4287) or the *FOBG_10856* (Fo47) gene and normalized to the *M. polymorpha* Mp*EF1a* gene. Error bars indicate SD (n = 6). Asterisks indicate statistical significance versus mock (one way ANOVA, Bonferroni’s multiple comparison test, p < 0.05).

A comparative scanning electron microscopy (SEM) analysis of the dorsal and ventral surfaces of inoculated *M. polymorpha* thalli inoculated with Fol4287 and Fo47 during early infection stages (3 dpi) revealed a similar pattern of hyphal growth on the thallus surface with penetration events occuring mainly between epidermal cells. Moreover, direct penetration of plant cells by invasive fungal hyphae was occasionally observed (Fig. 1B). Confocal microscopy of thalli inoculated with Fol4287 and Fo47 strains expressing the green fluorescent protein clover showed predominantly intercellular growth of *F. oxysporum* hyphae (Fig. 1C). The observed pattern of intercellular hyphal penetration and growth resembles that previously reported for Fol4287 on tomato roots (33). RT-qPCR detected the presence of fungal biomass in infected thalli at 6 dpi, in contrast to the uninoculated control (Fig. 1D). We conclude that *F. oxysporum* infects and colonizes the non-vascular plant *M. polymorpha* mainly through intercellular hyphal growth, similar to that observed in the root cortex of an angiosperm host.

#### *F. oxysporum* causes maceration and killing of *M. polymorpha* thallus tissue

*M. polymorpha* thalli inoculated with Fol4287 or Fo47 exhibited visible signs of tissue maceration and cell killing in the infected areas, suggesting the release of plant cell wall degrading enzymes by the fungus (Fig. 2A). Previous work established that exo- and endopolygalacturonases (PGs) are secreted by *F. oxysporum* during different stages of tomato plant infection and contribute to fungal virulence (34–36). During infection of *Marchantia* a marked upregulation of the transcript levels of *pg1* and *pg5* encoding the two major endoPGs and *pgx6* encoding the major exoPG of *F. oxysporum* was detected (Fig. 2B). In Fol4287, expression of the two endoPGs increased progressively to reach high levels at 3 dpi and then dropped at 7 dpi, while expression of the exoPG followed the same pattern at lower expression levels. These findings reflect the same trend previously observed in the angiosperm host tomato (36). By contrast, in Fo47 expression of all 3 PGs was highest at 1dpi and dropped markedly during subsequent time points. Interestingly, endoPG5 expression in Fol4287 was at least one order of magnitude higher than in Fo47 (Fig. 2B), which could explain the lower level of maceration caused by the endophytic strain.

**Fig. 2.**
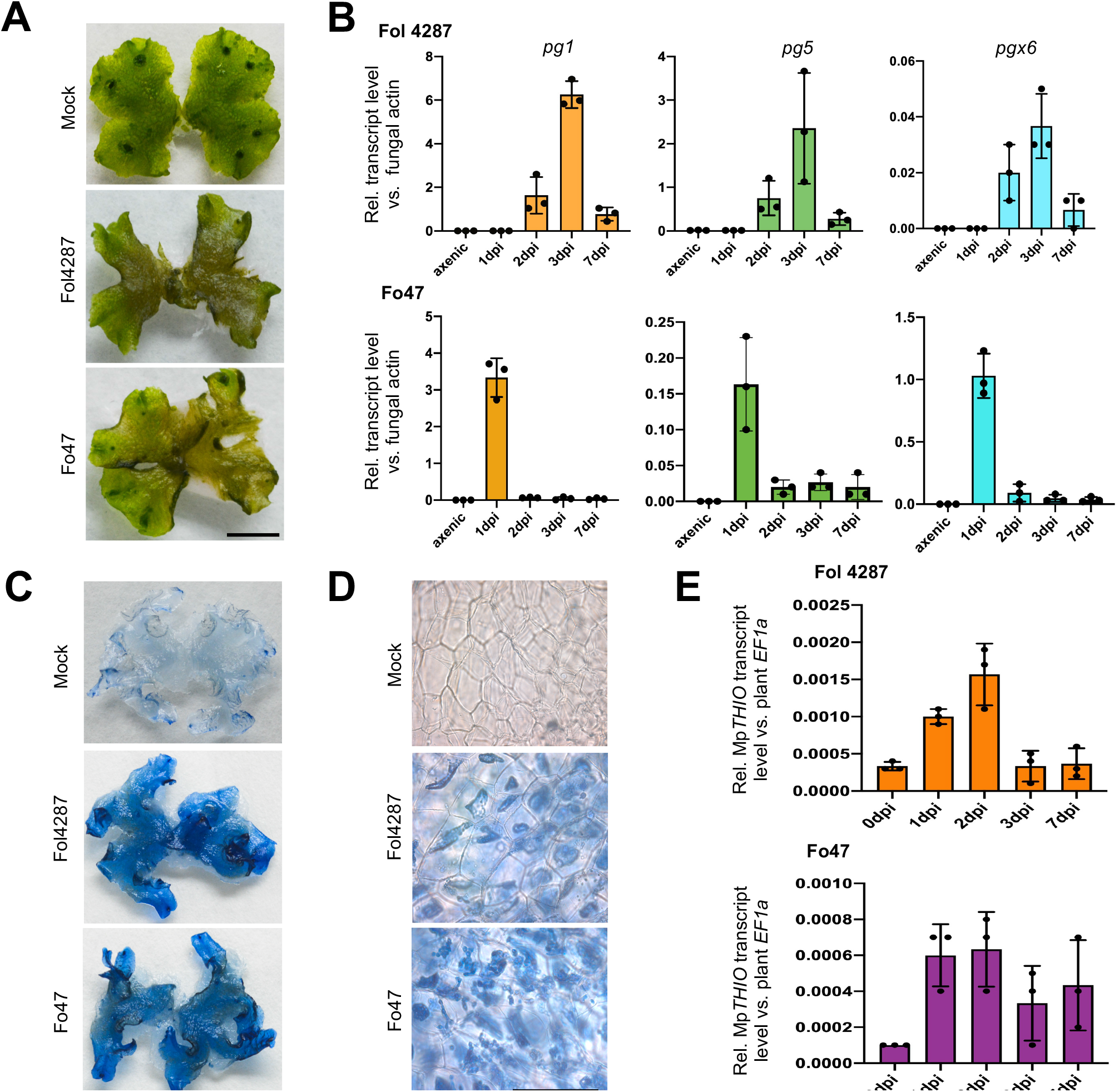
*F. oxysporum* strains cause plant tissue maceration and killing. **(A)** Images showing maceration of thallus tissue in *M. polymorpha* Tak-1 plants 7 days after dip inoculation with 5*10^6^ microconidia ml^−1^ of the indicated *Fo* strains or water (mock). Scale bar, 1 cm. **(B)** Transcript levels of the Fo genes *pg1*, *pg5* and *pgx6* encoding the two major endo- and the major exopolygalacturonases, respectively were measured by RT-qPCR of cDNA obtained from *M. polymorpha* Tak-1 plants at different times after dip inoculation with the indicated *Fo* strains or from fungal mycelium grown in liquid minimal medium (axenic). Transcript levels were calculated using the ΔΔCt method and normalized to those of the *Fo* actin gene. Error bars indicate SD (n = 3). (**C,D**) Macro-**(C)** and microscopic **(D)** images of thalli subjected to trypan blue staining for visualization of killed plant cells at 5 dpi. All images are representative of three independent experiments. Scale bar, 1 cm **(C)** or 100 μm **(D)**. **(E)** Transcript levels of the *M. polymorpha* cell death marker gene Mp *THIO* in *M. polymorpha* Tak-1 plants at different timepoints after dip inoculation with the indicated *Fo* strains were measured as described in **(B)** and normalized to those of the Mp*EF1a* gene. Error bars indicate SD (n = 3).

To confirm killing of *M. polymorpha* cells by *F. oxysporum* we performed Tryphan blue staining, which revealed the occurance of cell death in thalli infected with either Fol4287 or Fo47 at 5dpi, in contrast to the uninoculated control (Fig. 2C). Microscopy analysis confirmed staining of *M. polymorpha* cells, showing death signatures in the colonized thalli infected by Fol4287 and Fo47 (Fig. 2D). Moreover, the expression of the *M. polymorpha* cell death marker gene Mp*THIO* (Mapoly0001s0057) (26) was upregulated in thallus tissue infected with either Fol4287 or Fo47, with transcript levels in Fol4287 being approximately twice as high as those in Fo47 at earlier timepoints of 1 and 2dpi (Fig 2E). We conclude that the tissue collapse and maceration observed in infected *M. polymorpha* thalli is associated with secretion of plant cell wall degrading enzymes such as polygalacturonases and killing of host cells by *F. oxysporum*, which likely ensures nutrient support of the pathogen during host colonization. This was accompanied by production of fungal microconidia and the chlamydospores on the macerated thalli tissue which serve as structures for pathogen dispersal and survival (*SI Appendix*, Fig. S2A-B).

#### *M. polymorpha* senses *F. oxysporum* molecular patterns to activate the plant defense response

Plants have evolved conserved mechanisms to detect microbial invaders via pathogen associated molecular patterns (PAMPs) that trigger an efficient immune response (3–4). Here we asked whether *M. polymorpha* can perceive molecular patterns from *F. oxysporum*, as previously shown for PAMPs from *P. syringae* (26). Addition of crude boiled extracts from Fol4287 or Fo47 mycelia to *Marchantia* Tak-1 gemmalings grown in liquid media induced a concentration dependent growth inhibition (Fig. 3A and 3B, *SI Appendix*, Fig. S3A). Moreover, in contrast to the mock control, *M. polymorpha* thalli treated with *F. oxysporum* extracts showed rapid upregulation of orthologs of *Arabidopsis thaliana* PAMP responsive genes *CML42* (Mapoly0038s0010) and *WRKY22* (Mapoly0051s0057) (26, 37) (*SI Appendix*, Fig. S3B). These results suggest that crude extracts from Fol4287 and Fo47 contain molecular patterns that trigger a PAMP perception response in *M. polymorpha*.

**Fig. 3.**
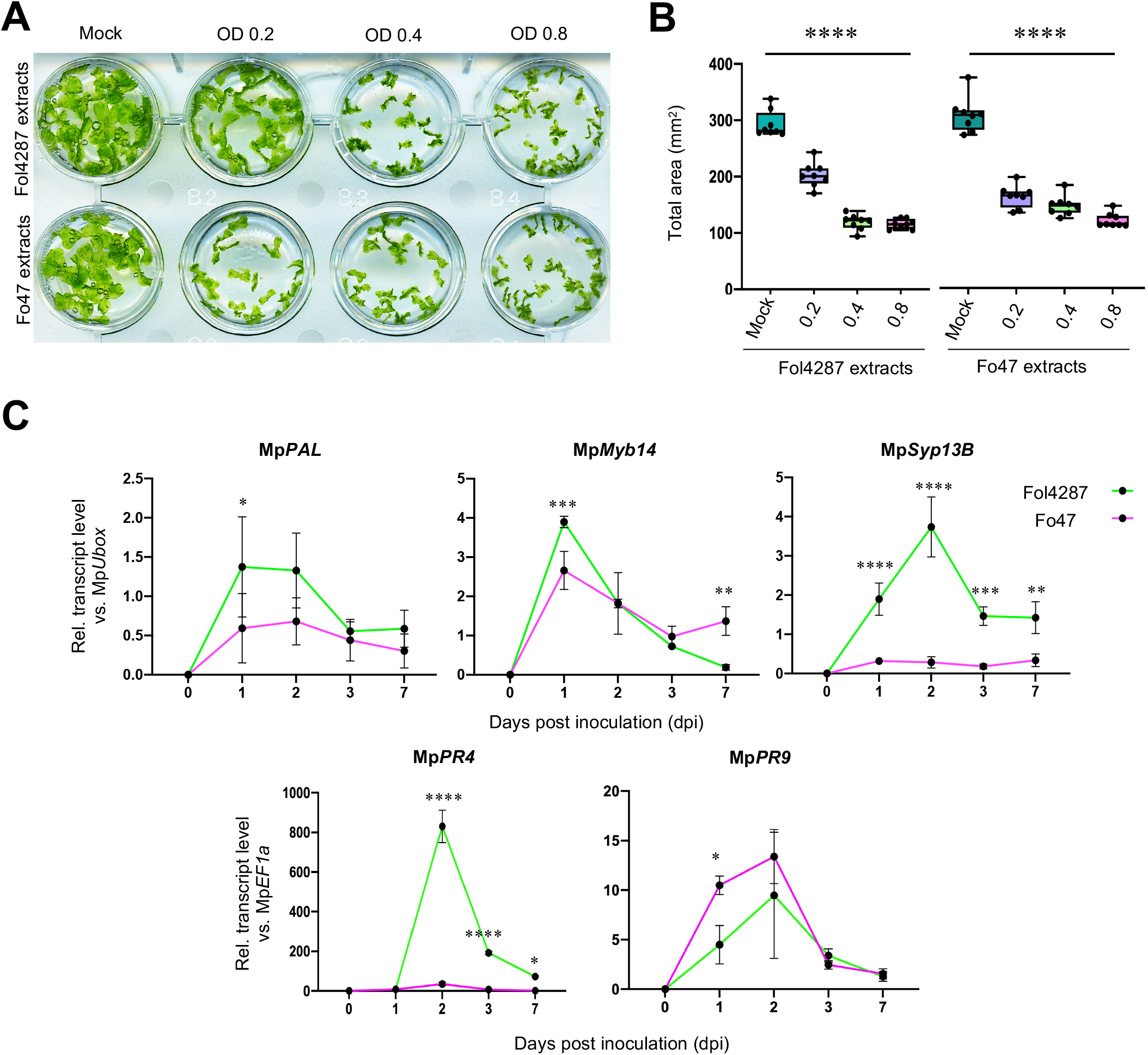
*M. polymorpha* perceives pathogen-associated molecular pattern (PAMP) signatures from *F. oxysporum* and induces a plant defense response. **(A-B)** Growth inhibition of *M. polymorpha* in response to *F. oxysporum* PAMPs. Tak-1 (male) gemmalings were grown for 14 days in liquid medium containing different concentrations of crude boiled extracts from Fol4287 or Fo47 (OD600 = 0.2, 0.4, 0.8) or water (Mock). **(A)** Images of gemmalings. **(B)** Quantification of total area (mm^2^) of gemmalings from **(A)**. Error bars represent SD (n = 8). Statistical significance compared to mock-treated plants (p < 0.01) is indicated by an asterisk. **(C)** *M. polymorpha* mounts a defense response in response to infection by *F. oxysporum.* Transcript levels of defense-reklated genes Mp*PAL* (phenylalanine ammonia lyase), Mp*Myb14 (*transcription factor), Mp*Syp13B* (membrane syntaxin) Mp*PR4* (chitin binding protein) and Mp*PR9* (peroxidase) were measured by RT-qPCR of cDNA obtained from *M. polymorpha* Tak-1 plants 1, 2, 3 and 7 dpi after dip inoculation with the indicated fungal strain. Transcript levels for each sample were calculated using the ΔΔCt method and normalized to those of the Mp*EF1a* or Mp*U-box* gene. Error bars indicate SD (n = 3). Different stars indicate statistically significant differences in transcript abundance between the two strains tested (2 way ANOVA, Bonferroni’s multiple comparison test, p < 0.05). Error bars represent SD.

Next, we wondered whether *M. polymorpha* activates a defense response upon infection by *F. oxysporum*, as previously shown for bacterial and oomycete pathogens (26, 29). A marked upregulation of genes encoding the flavonoid biosynthesis components Mp*PAL* and Mp*Myb14* as well as the membrane syntaxin Mp*SYP13B* was detected in thalli inoculated with Fol4287 or Fo47 (Fig. 3C). A similar response was observed for the pathogenesis-related proteins Mp*PR4* (chitin-binding) and Mp*PR9* (peroxidase), with Mp*PR4* showing a much higher induction in response to Fol4287. Taken together these data suggest that *M. polymorpha* senses PAMPs from different *F. oxysporum* strains and mounts a characteristic defense response upon fungal infection.

#### Core pathogenicity mechanisms, but not host-specific virulence effectors are required for *F. oxysporum* infection on *M. polymorpha*

To infect plants, fungal pathogens have evolved ancient and broadly conserved pathogenicity mechanisms as well as more recent, host specific virulence effectors. Previous studies in *F. oxysporum* identified a number of core pathogenicity factors including two mitogen activated protein kinases (MAPKs) Fmk1 and Mpk1, which control invasive growth and cell wall integrity, respectively (38, 39), the β-1,3-glucanosyltransferase Gas1 involved in cell wall assembly (40) or the zinc finger transcription factor Con7-1, which regulates hyphal morphogenesis and infection (41). Here we found that isogenic Δ*fmk1,* Δ*mpk1,* Δ*gas1* and Δ*con7-1* mutants of Fol4287 caused reduced disease symptoms and accumulated significantly less fungal biomass in *M. polymorpha* thalli than the wild type strain (Fig. 4A and 4B). These results reflect the same trend as those obtained in tomato plants, including a significant reduction in wilt symptoms, mortality and fungal biomass in infected roots and stems (Fig. 4C-E, *SI Appendix*, Fig. S4A-B).

**Fig. 4.**
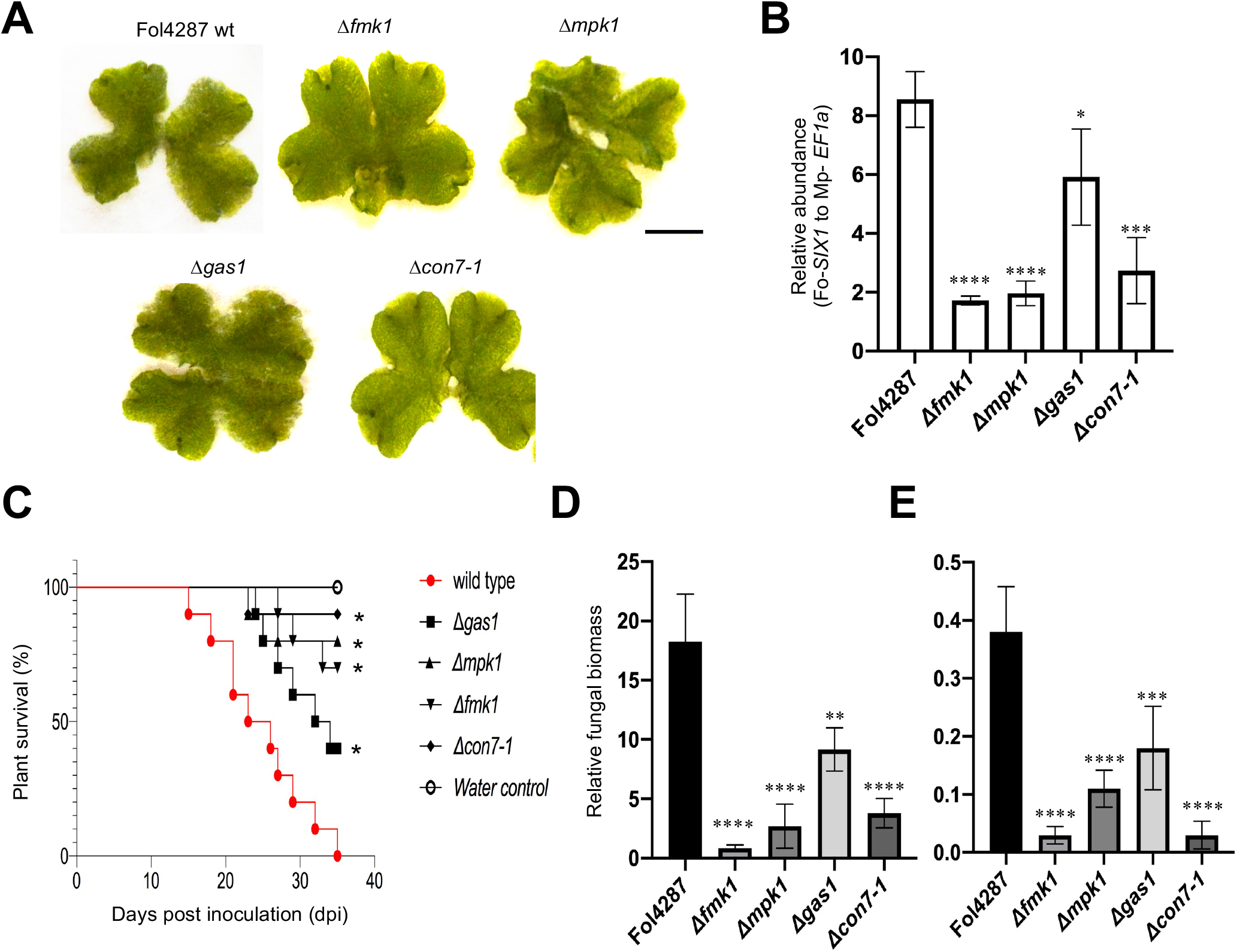
Core pathogenicity mechanisms are required for *F. oxysporum* infection on *M. polymorpha*. **(A)** Macroscopic disease symptoms on *M. polymorpha* Tak-1 plants 5 days after dip inoculation with 10^5^ microconidia ml^−1^ of the Fol4287 wild type strain (wt) and isogenic mutants in the indicated genes. Images are representative of three independent experiments. Scale bar, 1 cm. **(B)** Fungal burden determined 6 days after inoculation in infected thalli. The relative amount of fungal DNA quantified with Fo-*SIX1* was normalized to the Mp*EF1a* and expressed relative to the wild type (wt) (*P < 0.05, versus wild type according to unpaired t-test). Error bars indicate SD; n = 3. **(C)** The tested core pathogenicity determinants are required for full virulence. The Kaplan–Meier plot shows survival of tomato plants infected with Fol4287. Number of independent experiments (*n_i.ex_*.) = 3; 10 plants/treatment. Data shown are from one representative experiment. * *P* < 0.05 versus Fol4287 alone according to log-rank test. **(D, E)** Quantification of fungal biomass in roots **(D)** or stems **(E)** of tomato plants 10 days after dip inoculation with the indicated Fo strains or water (mock). Fungal biomass was measured by RT-qPCR using specific primers for the Fo4287 *ppi* gene, normalized to the tomato *Sl-GADPH* gene and expressed relative to the Fo4287 wt in roots and stem. Statistical significance versus wt (p < 0.05, one way ANOVA, Bonferroni’s multiple comparison test) is indicated by an asterisk. Error bars indicate SD (n = 3).

Host-specific pathogenicity of Fol4287 on tomato plants is conferred by the LS chromosome 14, which encodes a suite of Six effectors (17). Loss of this chromosome leads to inability to cause vascular wilt on tomato, while deletion of individual effector genes such as *six1* or *six3* genes causes reduced virulence (19, 42, 43). Here we found that Δ*six1* and Δ*six3* mutants showed no detectable differences in disease symptom severity caused on Tak-1 thalli as compared to Fol4287 (Fig. 5A). Moreover, upregulation of *six1* and *six3* transcript levels in Fol4287 was increased by 4 and 3 orders of magnitude, respectively, during infection of tomato roots as compared to *M. polymorpha* (Fig. 5B). We conclude that core pathogenicity mechanisms of *F. oxysporum* are largely conserved between tomato and liverwort, whereas Six effectors encoded by LS regions are dispensable for infection of *M. polymorpha*.

**Fig. 5.**
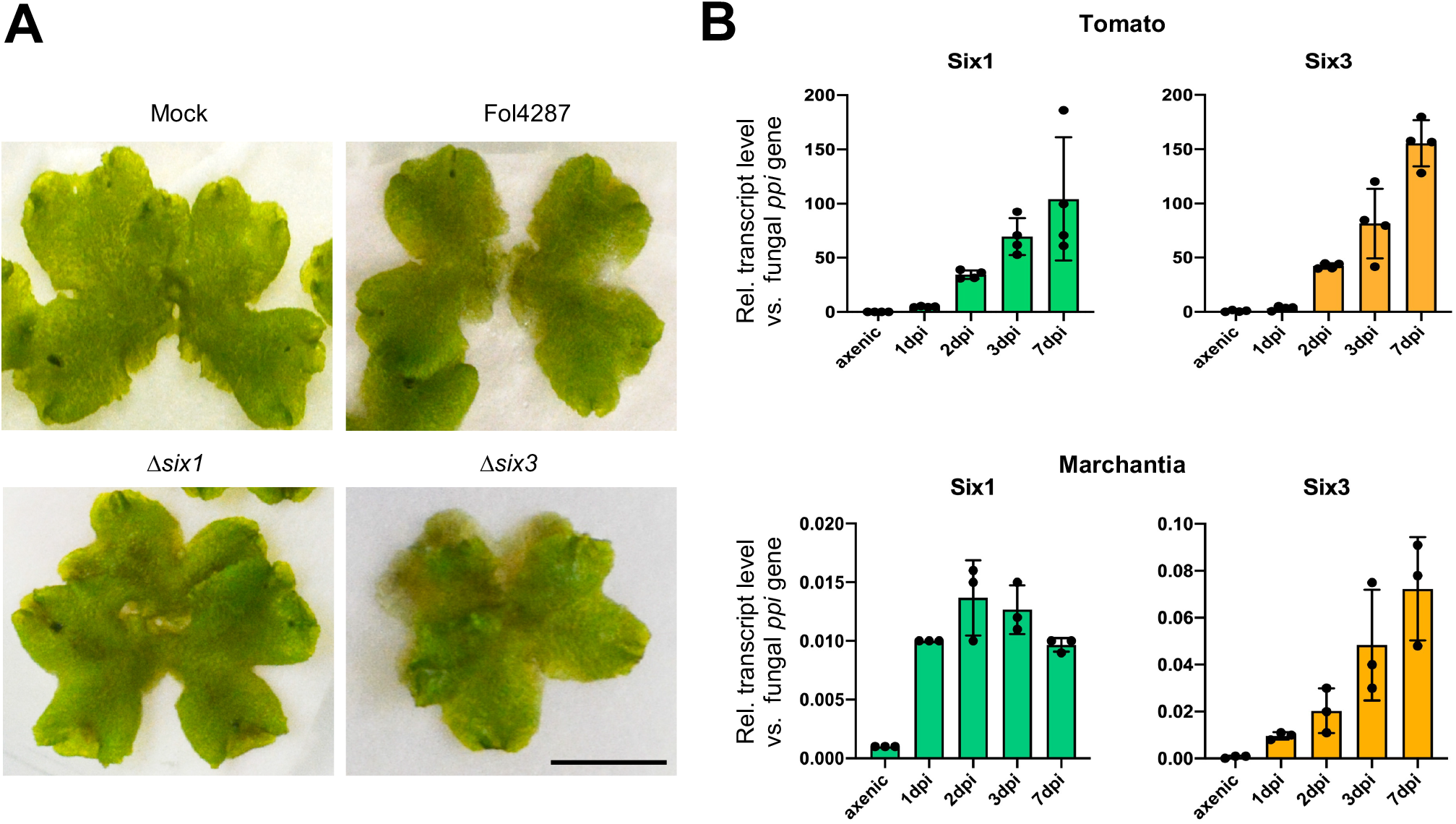
Tomato host-specific Six effectors are dispensable for *F. oxysporum* infection on *M. polymorpha.* **(A)** Macroscopic disease symptoms on *M. polymorpha* Tak-1 plants 5 days after dip inoculation with 10^5^ microconidia ml^−1^ of the Fol4287 wild type strain (wt) and mutants in the indicated genes. Images are representative of three independent experiments. Scale bar, 1 cm. **(B)** Transcript levels of the Fo4287 genes *six1* and *six3* encoding lineage specific (LS) effectors secreted in xylem (SIX) were measured by RT-qPCR of cDNA obtained from tomato roots (upper) or *M. polymorpha* Tak-1 plants (lower) 1, 2, 3 and 7 dpi after dip inoculation with Fol4287 or from fungal mycelium grown in liquid minimal medium (axenic). Transcript levels for each sample were calculated using the ΔΔCt method and normalized to those of the *Fo-ppi* gene. Error bars indicate SD (n = 3). Note that *in planta* upregulation of *six* effector genes is approximately four orders of magnitude higher in the angiosperm host Tomato compared to the liverwort *Marchantia*.

#### *F. oxysporum* induces RALF-independent alkalinization in *M. polymorpha*

During infection, fungal pathogens often induce extracellular alkalinization to promote host colonization (44). Fol4287 was previously shown to secrete a functional homologue of the plant regulatory peptide Rapid Alkalinizing Factor (RALF) which triggers alkalinization of the host apoplast and increases virulence on tomato plants (16). Using a plate bioassay with the pH indicator bromocresol purple, we found that Fol4287 and Fo47 triggered a marked extracellular alkalinization around *M. polymorpha* thalli, in contrast to only a slight alkalinization detected in presence of the fungus alone (*SI Appendix*, Fig. S5A). Unexpectedly, the Fol4287 Δ*ralf* mutant and a strain overexpressing *ralf* (16) induced a similar alkalinization response and caused disease symptoms that were indistinguishable from those of the wild type strain (*SI Appendix*, Fig. S5B). We conclude that *F. oxysporum* induces alkalinization during infection of *M. polymorpha* through a RALF-independent mechanism.

### Discussion

The availability of the *M. polymorpha* genome and the establishment of biotic interaction systems with oomycete and bacterial pathogens (26, 28–30) has provided new opportunities to explore plant-pathogen co-evolution across evolutionary timescales (2, 24–25). Compared to the aerial plant parts, immune responses in roots have been explored in less detail and likely represent a more complex scenario, due to the continuous exposure to a plethora of both beneficial and pathogenic microbes which constitute the plant microbiome (45, 46). Here we report a robust experimental infection system in *M. polymorpha* based on *F. oxysporum,* an economically important broad host range fungal pathogen. We used the *Marchantia* model to identify conserved pathogenicity mechanisms of this root-infecting wilt fungus that are shared during infection of a vascular plant host and a bryophyte lacking true roots and vasculature. Furthermore, we compared infection and disease development between a tomato pathogenic (Fol4287) and an endophytic fungal isolate (Fo47).

The infection cycle of *F. oxysporum* in angiosperm hosts consists of 3 distinct phases (Fig. 6): 1) penetration of the root and asymptomatic intercellular growth in the cortex; 2) crossing of the endodermis, entry in the xylem vessels and systemic colonization of the host resulting in plant death; 3) extensive maceration of the moribund plant tissue and development of dispersal and resting structures (micro- and macroconidia, chlamydospores) (13). Phases 1 and 3 are controlled mainly by core pathogenicity factors (14, 33, 36, 38–41), whereas phase 2 critically depends on host specific effectors encoded on LS genomic regions (17–20, 42, 43). Here we investigated, the mode of infection of a vascular fungal pathogen on a non-vascular plant species. *F. oxysporum* efficiently colonized *M. polymorpha* thalli and provoked clearly visible disease symptoms similar to those previously reported in the oomycete pathogen *P. palmivora* (29). In contrast to the characteristic wilt disease observed on angiosperm hosts, infection of *F. oxysporum* on *M. polymorpha* resulted in more general disease symptoms such as tissue browning, maceration and cell death, likely caused by secretion of plant cell wall degrading enzymes such as polygalacturonases. Importantly, *F. oxysporum* was able to complete its infection cycle on the bryophye host including the production of conidia and chlamydospores, representing the main dispersal and survival structures of this pathogen during vascular wilt disease on angiosperms (15).

**Fig. 6.**
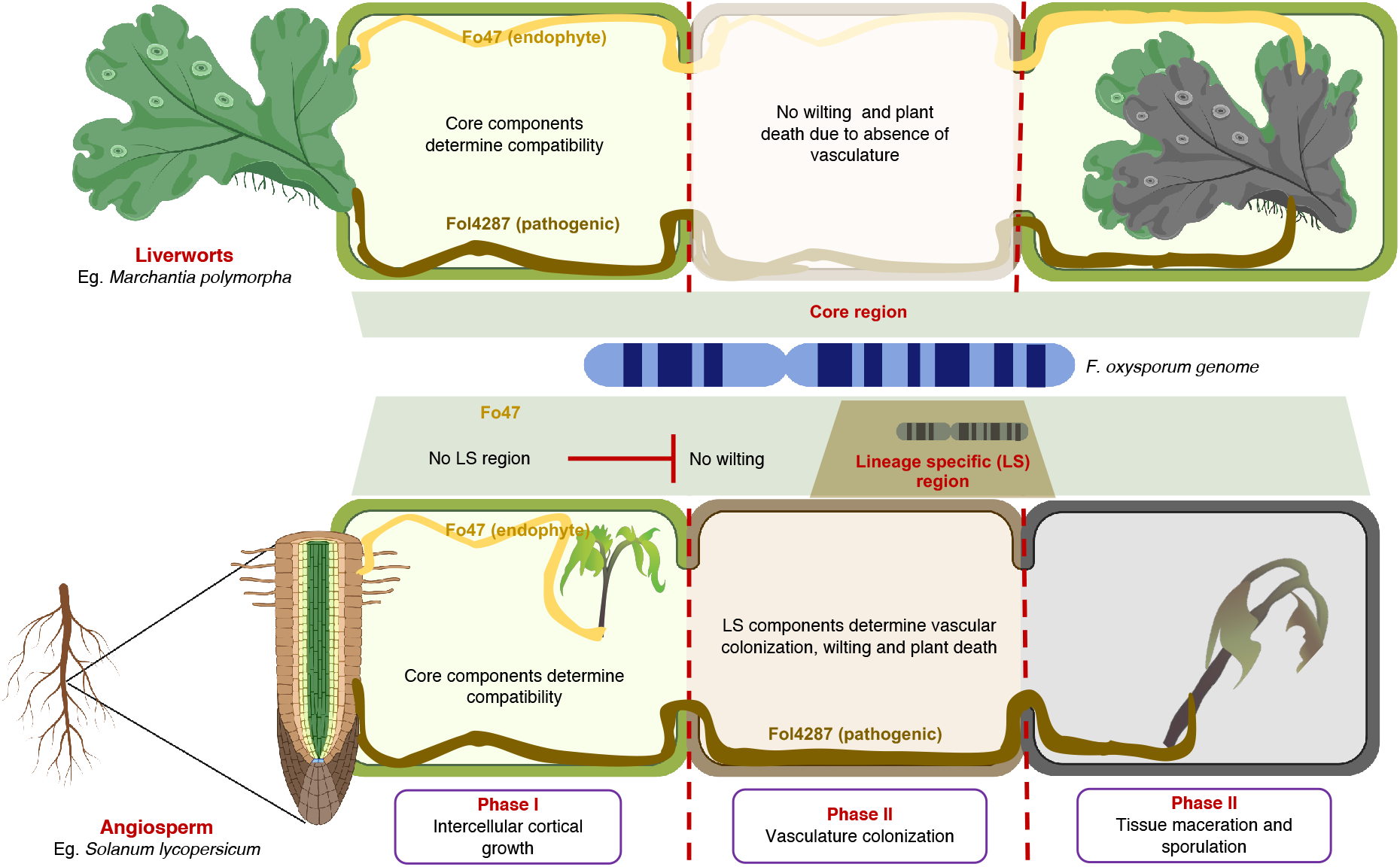
Distinct infection strategies of *F. oxysporum* on vascular and non-vascular plants. Schematic diagram illustrating distinct infection strategies employed by *F. oxysporum* during infection of a vascular (Tomato) and a non-vascular plant host (*M. polymorpha*). In angiosperm hosts, the infection cycle consists of 3 phases (lower panel): 1) asymptomatic intercellular growth in the root cortex; 2) entry in the xylem vessels and systemic colonization of the host resulting in plant death; 3) extensive maceration of the moribund plant tissue and development of dispersal and resting structures (micro- and macroconidia, chlamydospores). Phase 2 requires host specific effectors encoded on lineage specific (LS) genomic regions, whereas phases 1 and 3 depend mainly on pathogenicity factors encoded on core regions. Infection of the non-vascular host *Marchantia* lacks phase 2 due to the absence of a vasculature (upper panel) and thus depends exclusively on core pathogenicity factors. The model suggests that systemic colonization and wilting of the host plant by fungal pathogens evolved after the emergence of vascular land plants and that the core infection mechanisms may be ancient and likely evolved before the evolutionary divergence of non-vascular and vascular plants. Parts of this figure were created using Biorender. https://biorender.com/

Our results obtained with isogenic gene deletion mutants demonstrate that infection of *F. oxysporum* in *Marchantia* requires broadly conserved pathogenicity factors that are encoded on core genomic regions such as MAPKs, transcription factors and cell wall remodeling enzymes. Host alkalinization, another conserved pathogenicity mechanism (44), was also observed during infection of *Marchantia* by *F. oxysporum*. Intriguingly, in contrast to tomato (16), alkalinization in liverwort was independent of the fungal RALF peptide. This could be due to differences in the recognition profile of the plant receptor kinase FERONIA, which mediates alkalinization in response to both plant and fungal RALFs (16, 47). Collectively, our findings demonstrate that core infection mechanisms of *F. oxysporum* are conserved across evolutionarily distant host plant lineages, implying that these ancient fungal pathogenicity determinants predate the separation into vascular and non-vascular plants which occurred more than 450 million years ago (2).

The presence of core pathogenicity mechanisms controlling invasive hyphal growth and colonization of living plant tissue explains why an endophytic *F. oxysporum* isolate, which lacks host specific virulence effectors and fails to cause wilting on angiosperm hosts, is able to induce similar disease symptoms in this ancient plant lineage as a tomato pathogenic strain (48). In line with this idea, we found that the *six1* and *six3* genes, which are highly upregulated and contribute to virulence on tomato, are dispensable for infection of *M. polymorpha*. The differential role of LS effectors on angiosperm and bryophyte plants can be adscribed to the lack of a true vasculature in the liverwort, which results in absence of phase 2 of the disease cycle and thus prevents rapid host colonization via the xylem vessels (Fig. 6). The absence of systemic infection in *Marchantia* is further supported by the finding that, in spite of the severe disease symptoms observed at the centre of the thallus, all plants survived the challenge and resumed apical growth at the meristematic tissue.

An important goal in Evo-MPMI is to understand how plant immunity has evolved. We observed a marked growth inhibition of *M. polymorpha* upon exposure to *F. oxysporum* extracts, mirroring that previously observed with the bacterial pathogen *P. syringae* (26). These findings confirm that this non-vascular liverwort can sense PAMPs from different types of microbial pathogens and are further supported by the marked upregulation of plant defense markers such as syntaxin Mp*SYP13B* (29), the flavonoid biosynthesis components *PAL* and *Myb14b* or the pathogenesis related proteins *PR4* and *PR9*. Since the *M. polymorpha* genome only contains about a third of the receptor kinases found in *A. thaliana* (24), the availability of a rapid and robust PTI-like response to *F. oxysporum* extracts will facilitate the identification of novel fungal PAMPs and cognate PRRs that trigger immune activation in bryophytes.

In summary, our work on this newly established pathogen-host system reveals that *F. oxysporum,* a root-infecting vascular wilt fungus in angiosperms has a basal capacity to cause disease in non-vascular plants. Remarkably, two fungal strains with opposed lifestyles - pathogenic versus endophytic - similarly behave as pathogens in *Marchantia* because they share a common set of pathogenicity factors required for infection of both vascular and non-vascular plants. The lack of a role of host-specific virulence effectors, which are crucial for xylem colonization in angiosperm hosts, suggests that systemic wilt disease evolved in fungal pathogens after the emergence of vascular plants. These findings further highlight the potential of the *Marchantia* infection model to advance our understanding on the evolution of fungus-plant interactions.

### Materials and Methods

#### Strains and growth conditions

The tomato pathogenic isolate *F. oxysporum* f. sp. *lycopersici* 4287 (NRRL3436) and its derived mutants, the endophytic biocontrol isolate Fo47 (NRRL54002), as well as transformants of these strains expressing the GFP derivative clover were used throughout the study (*SI Appendix*, Table S1). For microconidia production, fungal strains were grown in potato dextrose broth (PDB) supplemented with the appropriate antibiotic(s) for 3 days at 28°C at 170 rpm. Microconidia were collected by filtration through monodur membranes and centrifugation as described (34) and counted in a Thoma chamber under a Olympus microscope.

#### *M. polymorpha* growth conditions and infection assay

*M. polymorpha* accessions Takaragaike-1 (Tak-1; male) was used. For *in vitro* assays, *M. polymorpha* gemmae were grown on plates of half Gamborg’s B5 medium (liquid medium) containing 1% agar (solid medium) at 21°C under a 16-h light/8-h dark cycle. For *F. oxysporum* infections, *M. polymorpha* thalli obtained from the cultures described above were grown for three weeks on plates of the same medium covered with a Whatman filter paper. Dip-inoculation was carried out by immersing the abaxial surface of the thalli for 30 minutes into a suspension of *F. oxysporum* microconidia at the desired concentration in a Petri dish together with the filter paper in order to cause minimum damage. For drop inoculation, a 5μl inoculum with desired spore concentration was applied on the thalli. Mock controls were treated with water. At least three thalli per treatment were used. Filter papers with the inoculated *M. polymorpha* thalli were transferred to a ray-sterilized microbox (Model Number. TP1600) fitted with an XXL+ filter (Labconsult, Brussels) containing vermiculite and incubated in a growth chamber at 21°C under short day conditions (10-h light/14-h dark). Disease symptoms were evaluated and imaged at the indicated days post inoculation (dpi). All infection experiments were performed at least three times with similar results, and representative images are shown.

#### Tomato root infection assays

Tomato root infection assays were performed as described (34). Briefly, roots of 2‐week‐old *Solanum lycopersicum* seedlings (cv. Monika) were immersed for 30 min in a suspension of 5 × 10^6^ microconidia ml^−1^ of the different strains and planted in minipots with vermiculite. Mock controls were treated with water. Ten plants per treatment were used. Plants were maintained in a growth chamber at 28°C under a 15-h light/9-h dark cycle. Plant survival was recorded daily. Mortality was calculated by the Kaplan−Meier method using GraphPad 9.0, USA.

#### Scanning electron microscopy

Sample preparation was carried out as reported (49) with minor modifications. Small pieces (1mm^2^) cut from infected *M. polymorpha* thalli at 3 dpi were fixed for 90 min with 2.5 % glutaraldehyde in 0.06 M Sorensen phosphate buffer at pH 7.2. After four washes in the same buffer for 10 min each, the samples were dehydrated in a graded series of increasing concentrations of ethanol (50 %, 70 %, 90 %, and 100 %) for 20 min per concentration, and critical point dried (Leica EM CPD 300; Leica Microsystems) using a customized program for plant leaves with a duration of 80 min (settings for CO2 inlet: speed=medium & delay=120s; settings for exchange: speed=5 & cycles=18; settings for gas release: heat=medium & speed=medium). Dried samples were mounted on aluminum stubs with carbon tape, sputter coated with 10 nm iridium (Leica EM ACE 600, Leica Microsystems) and imaged with a FEI Versa 3D scanning electron microscope (FEI, Hillsboro, OR, USA) under high vacuum conditions.

#### Generation of Fol-Clover or Fo47-Clover tagged *F. oxysporum* strains

Three copies in tandem of the *mClover3* gene (50) codon-optimized for *F. oxysporum* (*Fo-mClover3*) fused to a 3×FLAG tag coding sequence (3×*FLAG*) was cloned in the pUC57 plasmid backbone under control of the *Aspergillus nidulans gpdA* promoter and the *SV40* late polyadenylation signal. Fo-*mClover3*-labeled strains of Fol4287 and Fo47 (NRRL54002) were obtained by co-transforming protoplasts with the *Fo*-*mClover3* expression cassette and the hygromycin resistance cassette as previously described (51).

#### Laser scanning confocal microscopy

Laser Scanning confocal microscopy was performed using a Zeiss 880 Confocal microscope with Airyscan. *M. polymorpha* thalli inoculated with fluorescent transformants of Fol4287 or Fo47 expressing cytoplasmic 3× *mClover* were observed at an excitation of 488 nm and emission detected at 495-540 nm. To visualize plant cell walls, samples were co-stained by incubation in 2 mg ml^-1^ propidium iodide (PI) in water for 15 min in the dark as described (29). PI fluorescence was visualized at an excitation of 561 nm, and emission detected at 570-640 nm. Cells were visualized using the bright field DIC channel.

#### Quantification of fungal biomass and of gene expression *in planta*

For quantification of fungal biomass in tomato roots and stems or *M. polymorpha* thalli, the plant tissue was collected at the desired time points, snap frozen in liquid N_2_, finely ground to powder in a bead beater. Genomic DNA (gDNA) was extracted using a modified chloroform: octanol extraction protocol (52) and used for quantification of fungal biomass by real-time qPCR. Cycling conditions were 10 mins at 95°C followed by 40 cycles of 10 s at 95°C, 10 s at 62°C, and 20 s at 72°C. Data were analyzed using the double delta Ct method (53) by calculating the ratio of the plant housekeeping genes *SlGapdh* (tomato) or *MpEF1a* (*M. polymorpha*) versus the Fol4287-specific *six1* gene (*FOXG_16418*) or the Fo47-specific *FOBG_10856* gene. Primers used for qPCR analysis are listed in *SI Appendix*, Table S2.

#### Preparation of crude fungal extracts

Crude fungal extracts were prepared as described (26). Briefly, *F. oxysporum* cultures were grown in PDB for 3 days at 28°C and 170 rpm. The fungal mycelium was collected by filtration through a monodur membrane, resuspended in water at a ratio of 20% – 30% [fresh weight/volume], boiled in a water bath for 15 min at 95°C and cooled down to room temperature. The obtained crude extracts containing *F. oxysporum* PAMPs were stored at −20°C and used at an OD_600_ of 0.2-0.8 to evaluate the effect on *M. polymorpha* growth as described (26).

#### Analysis of gene expression by RT-qPCR

To measure transcript levels of fungal or plant genes in *M. polymorpha* or tomato, total RNA was isolated from snap frozen tissue of three biological replicates and used for reverse transcription quantitative PCR (RT-qPCR) analysis. Briefly, RNA was extracted using the Tripure Reagent and treated with DNAase (both from Roche). Reverse transcription was carried out with the cDNA Master Universal Transcriptor Mix (Roche), using 3μg of total RNA according to the manufacturer’s instruction. qPCR was performed using the CFX96 Touch™ Real-Time PCR Detection System (Bio-Rad). Primers used for qPCR analysis of different plant defense marker genes are listed in *SI Appendix*, Table S2. Data were analyzed using the double delta Ct method (53) by calculating the relative transcript level of the defense marker genes in relation to that of the house keeping reference genes Mp*U-box* or Mp*EF1a*. For expression analysis of fungal genes, the Fo peptidyl prolyl isomerase (*FOXG_08379*) or actin1 (*FOXG_01569*) genes were used as references.

For analysis of Mp*CML42* and Mp*WRKY22* gene expression, experiments were performed with RNA extracted from 7-to 14-day-old *M. polymorpha* grown on Gamborg’s B5 medium containing 1% agar. Typically, *M. polymorpha* was transferred to liquid Gamborg’s B5 medium two days prior induction with compounds. RNA extraction and cleanup was done using Trizol reagent (Invitrogen) followed by High Pure RNA Isolation Kit (Roche) and DNase digestion to remove genomic DNA contamination. First-strand cDNA was synthesized from 1 mg of RNA using the High Capacity cDNA Reverse Transcription Kit (Applied Biosystems), according to the manufacturers instructions. For quantitative PCR, five microliters from one-tenth diluted cDNA was used to amplify selected genes and the housekeeping gene Mp*U-box* using Power SYBR Green PCR Master Mix (Applied Biosystems). Primer sequences are described (*SI Appendix*, Table S2). Quantitative PCR was performed in 96-well optical plates in a 7500 Real Time PCR System (Applied Biosystems). Data analysis shown was done using three technical replicates from one biological sample. Error bars represent standard deviation (SD). In all cases, the measurements represent the ratio of expression levels between each sample and controls as indicated in each experiment. All samples were normalized against the housekeeping gene Mp*U-box*. All experiments were performed three times with similar results, and representative results are shown.

#### Whole Plant Alkalinization assays

Whole plant alkalinization assays on plates were performed as described previously with modifications (16). Briefly, 3-week-old *M. polymorpha* Tak-1 male thalli obtained as described above were placed on 0.5% water agar plates adjusted to pH 5.5 with 50% acetic acid and supplemented with 2μM of bromocresol purple, to which 1.25 × 10^6^ microconidia ml^−1^ of the appropriate *F. oxysporum* isolate or water (control) was added just prior to pouring. Plates were incubated in a growth chamber at 28°C under a 15-h light/9-h dark cycle. After 24 hours, the plates were imaged to document changes in pH.

## Supporting information

Suppmentary information

## Author Contributions

A.R., S.G.I., R.S. and A.D.P conceptualized and designed the research. A.R., S.G.I, M.S and B.Z conducted all experiments. A.R., S.G.I., M.S. and B.Z., performed the data analysis. A.R. and A.D.P. wrote the manuscript with input from all co-authors. All authors reviewed and approved the manuscript.

## Acknowledgements

A.R. and M.S. acknowledge funding from the European Union’s Horizon 2020 research and innovation program under the Marie Skłodowska‐Curie grant agreement No. 750669 and 797256. A.D.P. acknowledges support from the Spanish Ministry of Science and Innovation (MICINN, grant PID2019-108045RB-I00). A.R. acknowledges funding from Juan de la Cierva Incorporación grant from MICINN (IJC2018-038468-I). Work in the R.S. lab was funded by the Spanish Ministry of Science and Innovation grant PID2019-107012RB-I00/ AEI / 10.13039/501100011033. S.G.I. is funded by a Ramon y Cajal Fellowship RYC2019-026396-I, the CSIC grant 20212AT006 and the Spanish Ministry of Science and Innovation grant for young investigators RTI2018-094526-J-I00. We thank Martijn Rep for sending us the *F. oxysporum* Δ*six1* and Δ*six3* mutants and David Turrà for contributing the fluorescent *F. oxysporum* strains.

