## Supplementary material for "*Marchantia polymorpha* model reveals conserved infection mechanisms in the vascular wilt fungal pathogen *Fusarium oxysporum*": Suppmentary information

**This PDF file includes:**

**Figures S1 to S5**
**Tables S1 and S2**

### Fig. S1

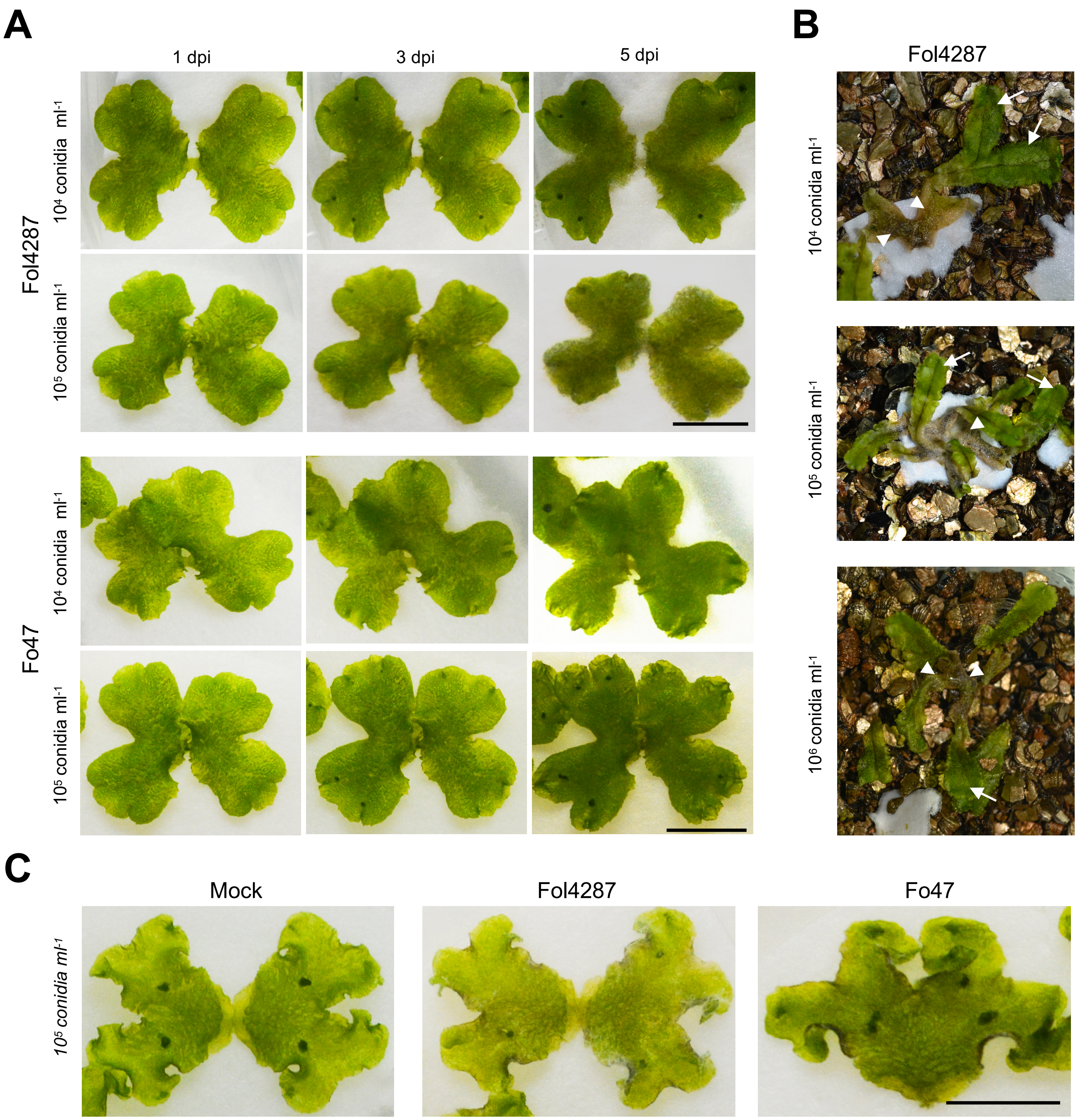

**Fig. S1. Effect of different inoculum concentrations and inoculation methods of *F. oxysporum* strains on thalli of *M. polymorpha***

**(A)** *M. polymorpha* Tak-1 plants 1, 3 and 5 days after dip inoculation with the indicated concentrations of microconidia of Fol4287 or Fo47. Scale bar, 1 cm.

**(B)** Meristematic areas comprising apical notches are more resistant to Fo infection than mature thalli. *M. polymorpha* Tak-1 plants 45 days after dip inoculation with the indicated concentrations of microconidia of Fol4287. Note that mature thalli are completely macerated (arrowheads) whereas young apical notches have escaped infection and are growing normally (arrows).

**(C)** *M. polymorpha* Tak-1 plants 5 days after drop inoculation with the indicated concentrations of microconidia of Fol4287 or Fo47 or water (Mock). Images are representative of three independent experiments. Scale bar, 1 cm.

### Fig. S2

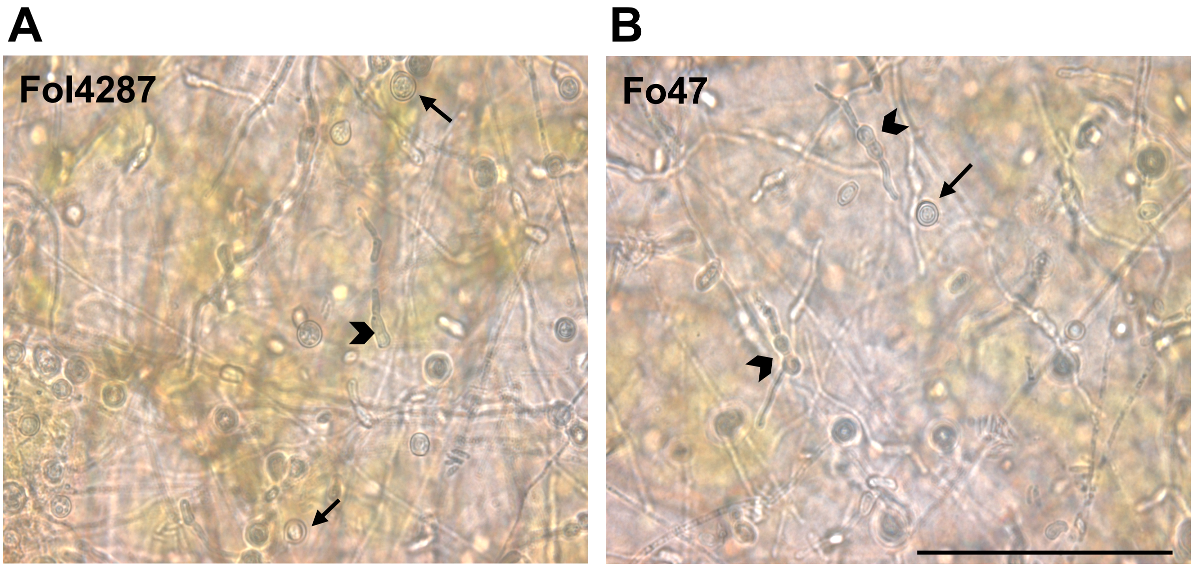

**Fig. S2. The macerated *M. polymorpha* shows sporulation of *F. oxysporum***
*M. polymorpha* Tak-1 plants 25 days after infection and complete maceration shows the production of microconidia (indicated by arrowheads) and chlamydospores (indicated by arrows) by both pathogenic (Fol4287) **(A)** and endophytic (Fo47) **(B)** strains of *F. oxysporum*. Scale bar, 100 µm.

#### Fig. S3

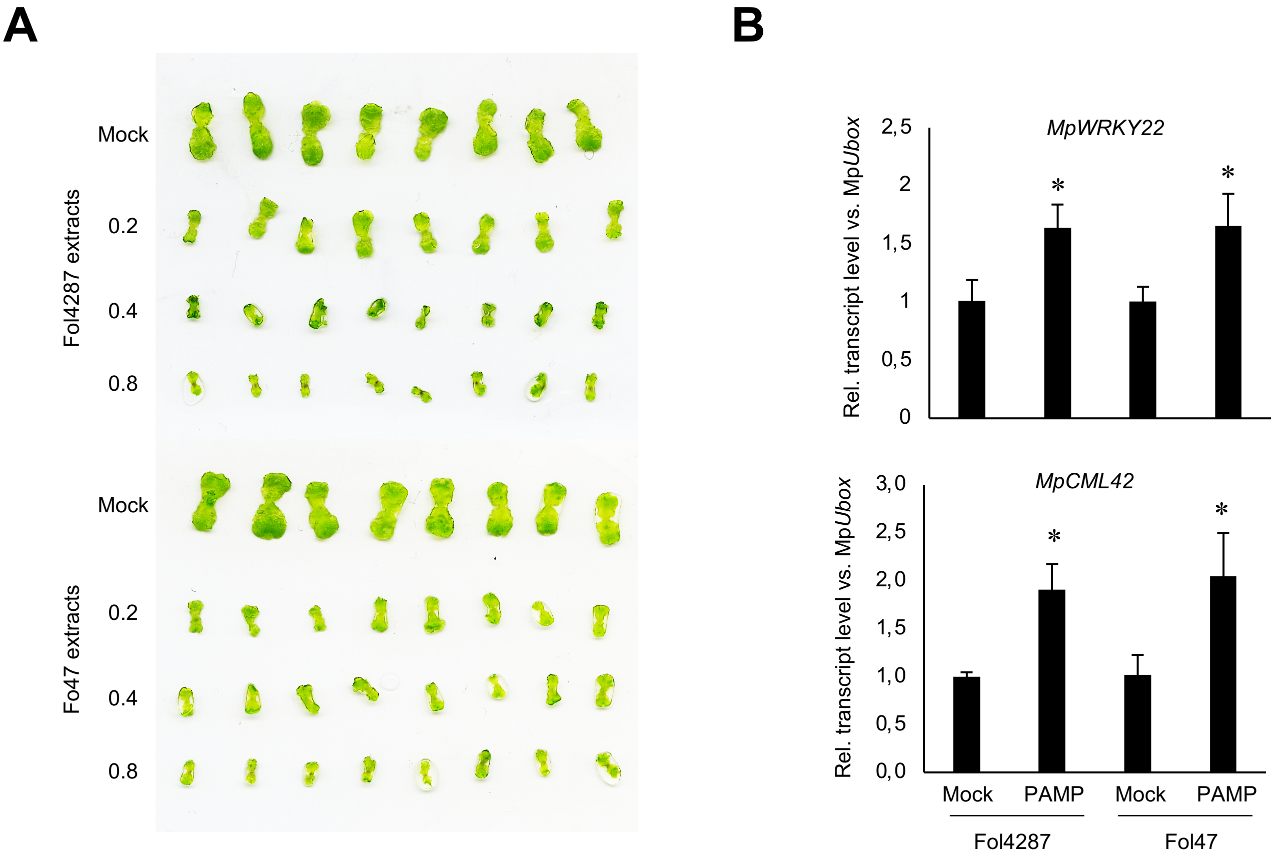

**Fig. S3. *M. polymorpha* responds to pathogen-associated molecular pattern (PAMP) signatures from different *F. oxysporum* strains**

**(A)** *M. polymorpha* Tak-1 (male) gemmalings were grown for 14 days in liquid medium containing different concentrations of crude boiled extracts from Fol4287 or Fo47 containing PAMP signatures (OD600 = 0.2, 0.4, 0.8) or water (Mock). Images of individual gemmalings are shown.
**(B)** Transcript levels of the *M. polymorpha* PAMP-responsive genes Mp*WRKY22* (upper) and Mp*CML42* (lower) were measured by RT-qPCR of cDNA obtained from *M. polymorpha* Tak-1 gemmalings 1 h after addition of crude boiled extracts from Fol4287 or Fo47 containing PAMP signatures (OD600 = 0.8) or water (Mock). Transcript levels for each sample were calculated using the ΔΔCt method and normalized to those of the Mp*Ubox* gene. Error bars indicate SD (n = 3). Statistical significance versus mock-treated plants ($p < 0.01$) is indicated by an asterisk.

Fig. S4

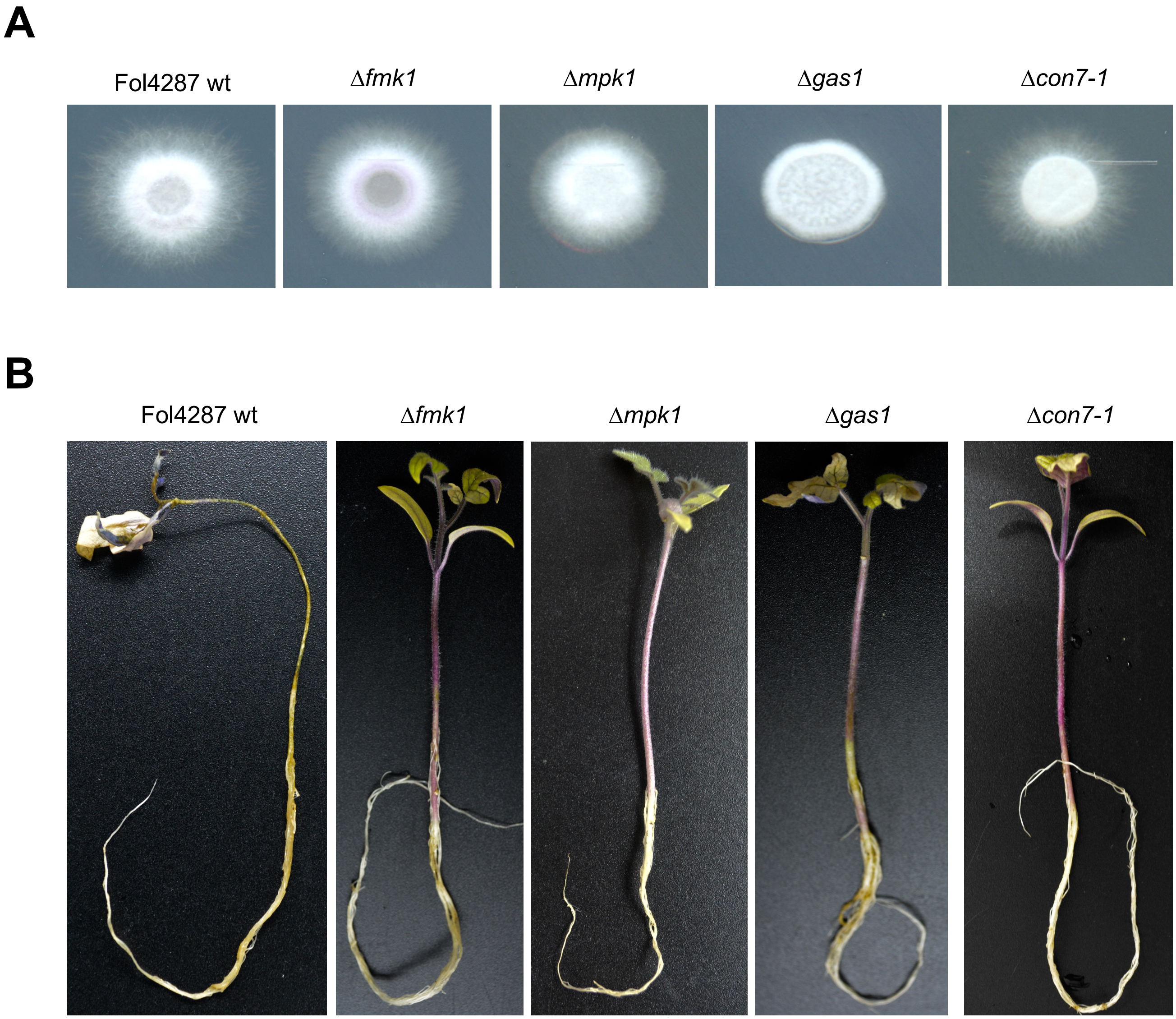

**Fig. S4. Role of different core pathogenicity determinants in colony development and virulence of Fol4287 on tomato plants.**

**(A)** Colony growth of the Fol4287 wild type strain (wt) and isogenic mutants in the indicated genes after 3 days on potato dextrose agar (PDA) plates.
**(B)** Representative images of tomato plants 25 days after root dip inoculation with microconidia of the wt and isogenic mutants in the indicated genes.

### Fig. S5

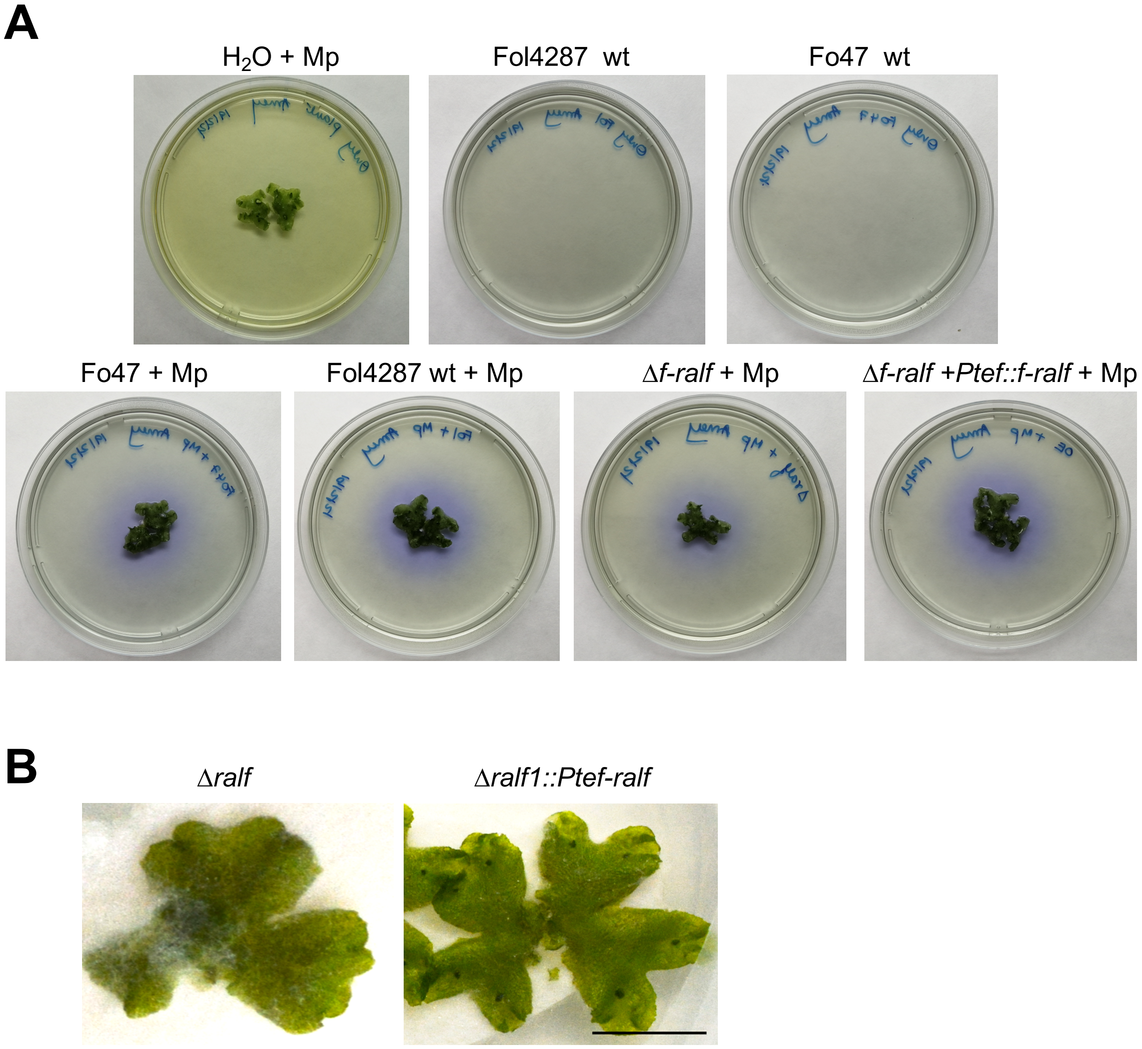

**Fig. S5. *F. oxysporum* induces F-RALF-independent alkalinization in *M. polymorpha*.**

**(A)** Plate alkalinization assay with *M. polymorpha*. Tak-1 thalli were placed on water agar plates adjusted to pH of 5.5 containing $1.25*10^6$ microconidia $ml^{-1}$ of the indicated *F. oxysporum* wild type strains (wt), the Δ*f-ralf* null mutant or the Δ*f-ralf* +*Ptef::f-ralf* overexpression strain or water as a control ($H_2O$) and 2 µM of the pH indicator bromocresol purple. Plates were imaged after 24 hours. Bright yellow, pH < 5.2; purple, pH > 6.8. Images are representative of three independent experiments. **(B)** *M. polymorpha* Tak-1 plants 5 days after dip inoculation with the indicated fungal strains. Images are representative of three independent experiments. Scale bar, 1 cm.

**Table S1. *Fusarium oxysporum* strains used in the study**

| Strain | Genotype | Gene Function | Reference |
|---|---|---|---|
| NRRL34936 (4287) | *Fusarium oxysporum* f.sp. *lycopersici* (race 2) - wildtype | | Ma et al., 2010 |
| NRRL54002 (Fo47) | *Fusarium oxysporum* Fo47 (biocontrol strain) | | Albouvette. 1999; Wang et al., 2020 |
| Fol4287-*mClover3* | *Fol4287::An-gpdaP:*3xFo-*mClover*3:3xFLAG | | This study |
| Fo47-*mClover*3 | *Fo47::An-gpdaP:*3xFo-*mClover*3:3xFLAG | | This study |
| *fmk1*Δ | *fmk1:: PHLEO* | MAPK | Di Pietro et al., 2001 |
| *mpk1*Δ | *mpk1:: HYG* | MAPK | Segorbe et al., 2017 |
| *gas1*Δ | *gas1:: HYG* | β-1,3-glucanosyltransferase | Caracuel et al., 2005 |
| *con7-1*Δ | *con7-1:: HYG* | Zinc finger transcription factor | Ruiz- Roldan et al., 2015 |
| *ralf*Δ | *f-ralf:: HYG* | rapid alkalinization factor | Masachis et al., 2016 |
| *ralf*Δ +Ptef-*f-ralf* | *f-ralf:: HYG; Ptef-ralf::PHLEO* | rapid alkalinization factor | Masachis et al., 2016 |
| *six1*Δ (FP822) | *six1::HYG* | effector | Rep et al., 2004 |
| *six3*Δ (FP1414) | *six3::HYG* | effector | Houterman et al., 2009 |

**Table S2. Oligonucleotide used in the study**

| Primer | Sequence | Use | Reference |
|---|---|---|---|
| MpU-box-Fwd /Rv | GAACTCCATGGCTTCTCTTT / GATGGTACAAGCCACAGGTA | Real time qPCR | Gimenez-Ibanez et al., 2019 |
| MpWRKY22-Fwd /Rv | AGATGCAGTAGCTCGAAAGG /CTTGAGATTGGCAGTTTCCT | Real time qPCR | Gimenez-Ibanez et al., 2019 |
| MpCML42-Fwd /Rv | GTTCCTCTCGCTCTACAGGT /TCATTCTCTCGCAGTTCTTC | Real time qPCR | Gimenez-Ibanez et al., 2019 |
| MpTHIO-Fwd /Rv | TCTGTCCATGGTGCTTTATC / ATAGACTGCACACGCGTACT | Real time qPCR | Gimenez-Ibanez et al., 2019 |
| MpEF1a-Fwd /Rv | CCGAGATCCTGACCAAGG /GAGGTGGGTACTCAGCGAAG | Real time qPCR | Carella et al., 2018 |
| MpPAL-Fwd /Rv | AATTCGCTGGGGCTCATTTC /ACAGAGCGCAACCATGAAAG | Real time qPCR | Kubo et al., 2018 |
| MpPRX (PR9)-Fw /Rv | ATTTCGATTGCTTCCGAGGC /AATCCCACATCCCCGAAGTT | Real time qPCR | Carella et al., 2018 |
| MpPR4-Fwd /Rv | TTCTGTGGTTTGCAGTTTCTTG /CGCCATTGTAGTGATTCGTTAG | Real time qPCR | Carella et al., 2019 |
| MpSYP13B-Fwd/Rv | CGAAATGAGCACGAACGGGA /CCAGCTTCTTCAGCAGCGAC | Real time qPCR | Carella et al., 2018 |
| MpMyb14-Fwd /Rv | TCGAAACTCTTCCACAGACAGA / GCTAATGAAGCCCGTACATAGG | Real time qPCR | Carella et al., 2019 |
| Fo-actin Fwd /Rv | ATGTCACCACCTTCAACTCCA / CTCTCGTCGTACTCCTGCTT | Real time qPCR | Bravo Ruiz et al., 2016 |

| | | | |
|---|---|---|---|
| Fopg1 Fwd/ Rv | GTCACTTCGGGTACAAACATC / CCTTGATGAACTTGATGCCGC | Real time qPCR | Bravo Ruiz et al., 2016 |
| Fopg5 Fwd /Rv | GCCTGGTCGCCTCCGTACT / TCTTCTTGCCGCCGTTGCTGCCCTTGCCGT | Real time qPCR | Bravo Ruiz et al., 2016 |
| Fopgx6 Fwd /Rv | GAAGTCATCGCAAGGTCTATAC / AGAACAGAATAGGTCGGAGGTA | Real time qPCR | Bravo Ruiz et al., 2016 |
| Fo-ppi Fwd /Rv | AAGGGTGACCAGTTCGATAG / TTCTCGCCGAGCTTCATTTG | Real time qPCR | This study |
| Fol4287_Six1- Fwd /Rv | GATCAGTGACCAGAGCTTGC / CCAGGCGATTTAGGCGATTC | Real time qPCR | This study |
| Fol4287_Six3- Fwd /Rv | CGAGCTTCAGCACCGAACCT / CGATCTCGTAGTTGGCGATG | Real time qPCR | This study |
| Fo47_10856_Fwd/ Rv | GTTTGGTTGGTTTGGCTCCC / GCAAGTCGTTCCTGCCCTTT | Real time qPCR | This study |
